# Experimental confirmation that an uncommon, yet clinically relevant mutation (G878A) in the *rrs* gene of *Mycobacterium tuberculosis* confers resistance to streptomycin

**DOI:** 10.1101/2021.09.14.460364

**Authors:** Pilar Domenech, Esma Mouhoub, Michael B. Reed

## Abstract

The effective treatment of patients diagnosed with drug resistant tuberculosis (TB) is highly dependent upon the ability to rapidly and accurately determine the antibiotic resistance/ susceptibility profile of the *Mycobacterium tuberculosis* isolate(s) involved. Thus, as more and more clinical microbiology laboratories advance towards the routine use of DNA sequence-based diagnostics, it is imperative that their predictive functions extend beyond the well-known resistance-conferring mutations, in order to also encompass as many of the lower-frequency mutations as possible. However, in most cases, the fundamental experimental proof that links these uncommon mutations with phenotypic resistance is still lacking. One such example is the G878A polymorphism within the *rrs* gene encoding the 16s rRNA. We, and others, have identified this mutation within a small number of drug-resistant *M. tuberculosis* isolates, although prior to this study a consensus regarding exactly which aminoglycoside antibiotic(s) it conferred resistance toward seems not to have been reached. Here we have employed oligo-mediated recombineering to specifically introduce the G878A polymorphism into the *rrs* gene of *M. bovis* BCG - a species very closely related to *M. tuberculosis* - and demonstrate that it confers low-level resistance to streptomycin alone. In our hands, it does not confer cross-resistance towards amikacin, capreomycin, nor kanamycin. We also demonstrate that the *rrs*^G878A^ mutation exerts a substantial fitness defect *in vitro*, that may at least in part explain why clinical *M. tuberculosis* isolates bearing this mutation appear to be quite rare. Overall, this study provides clarity to the resistance phenotype attributable to the *rrs*^G878A^ mutation and is relevant to the future implementation of genomics-based diagnostics, as well as the clinical management of patients in situations where this particular polymorphism is encountered.

## INTRODUCTION

Despite the availability of effective antibiotic therapy for close to 80 years, human tuberculosis (TB) resulting from infection with *Mycobacterium tuberculosis* remains as one of the leading causes of mortality in low-income countries worldwide (1). The very first anti-TB drugs, *para*-aminosalicylic acid (PAS) and streptomycin (STR), were first used to treat TB patients in 1944 and remain in use to this day for the treatment of multiply antibiotic-resistant TB under specific circumstances (2). Indeed, the evolution of multi- and extensively drug resistant (MDR and XDR, respectively) TB severely limits the effectiveness of current treatment programs, with close to 500,000 new MDR cases reported each year (3). Thus, as well as being the leading cause of death due to a single bacterial infection, *Mtb* also has the unenviable distinction of causing the most antimicrobial resistance-related deaths.

Treatment of patients with MDR and XDR-TB is extremely difficult and prohibitively costly, requiring multiple, potentially toxic drugs for up to 24 months (including some by injection), and is a major factor contributing to the ongoing global TB epidemic (3). In concert with the need for new and improved treatment regimens for these patients, is the need for the development of appropriate molecular diagnostic tools that can rapidly and accurately identify the specific resistance profiles of the *Mtb* isolates involved. It is imperative that this is achieved in order that the most effective antibiotic combinations can be administered as soon as possible after an initial diagnosis of TB is received.

Whilst a great technological advance, current culture-independent molecular tests (*i.e*. genotypic tests) for identifying antibiotic resistance in *Mtb*, including the PCR-based Gene Xpert^®^ and GenoType MTBDR systems, are limited to the identification of the most common of the known, or well-established, resistance-conferring mutations. In addition, depending upon the particular version of the system in use, they may also be limited to the detection of resistance to just 1 or 2 antibiotics (*e.g.* rifampicin (RIF) and/or isoniazid (INH)) (4–6). In contrast, next-generation sequencing (NGS) based approaches show tremendous potential for the unbiased detection of resistance mutations for DNAs prepared directly from patient sputum samples or from a primary culture. However, at present, NGS approaches are still somewhat limited by virtue of the fact that there remains many examples of low-frequency mutations or so-called “unexplained” resistance where poorly characterized mutations are detected that, whilst they may be suspected/ predicted to confer resistance or are associated with clinical resistance (with varying levels of confidence), have never actually been experimentally proven to confer phenotypic antibiotic resistance within a laboratory setting (7–11). The ability to confirm, or discount, as many of these lower-frequency mutations as possible would serve to increase the certainty with which antibiotic resistance and susceptibility predictions are able to be made based solely on genomic data. In turn, this would greatly enhance the clinician’s ability to correctly apply the most appropriate drug combinations as early as possible after a positive TB diagnosis is made, thus limiting treatment failure and further resistance development.

One example of an unconfirmed mutation was recently detected in our laboratory for a single MDR-TB isolate that is part of the McGill University/RI-MUHC strain collection comprised of 798 isolates collected over a six-year period from mostly foreign-born TB patients resident on the island of Montreal (12). As well as being resistant to RIF and INH, the strain was also classified as being resistant to STR by the Laboratoire de Santé Publique du Québec (LSPQ). As part of a separate ongoing study in our laboratory, we decided to identify the genetic basis for resistance to each of these antibiotics in this particular strain and sequenced the main candidate loci, including the *rpsL*, *rrs* and *gidB* genes, mutations in which are most frequently responsible for phenotypic STR resistance (10). Somewhat surprisingly, the only mutation detected in any of these gene sequences was a G to A SNP (single nucleotide polymorphism) at position 878 of *rrs* that encodes the 16sRNA sequence. More typical *rrs* mutations linked to STR resistance include the A514C and C517T SNPs (10, 13). Even more surprising was the fact that this mutation seems to be rarely recorded either in DNA sequence databases or the published literature. In fact, and as discussed below, in the small number of studies where this G878A SNP has been reported, there appeared to be some level of confusion regarding exactly which aminoglycoside antibiotics it may be associated with resistance towards - namely amikacin (AMK), capreomycin (CAP), kanamycin (KAN) and/or STR (14–18). As such, we considered it might be beneficial to the TB community for our group to determine experimentally whether or not this SNP contributes to resistance against one or more of these antibiotics currently in use for the treatment of drug resistant TB. Through the specific introduction of this point mutation into the antibiotic-sensitive *Mycobacterium bovis* Bacille Calmette-Guérin (BCG) background - a non-pathogenic member of the MTBC (*Mtb* complex) that is very closely related to *Mtb* at the genetic level - via oligo-mediated recombineering (recombination-mediated genetic engineering) (19, 20), herein we demonstrate for the first time that the G878A *rrs* mutation confers low-level resistance to STR alone.

## MATERIALS & METHODS

### Bacterial strains and culture

The MDR *Mtb* isolate #57001 was classified as belonging to the Euro-American lineage (Lineage 4) in a previous molecular epidemiological study of Montreal TB patient isolates collected between January 2001 and May 2007 (12). The *M. bovis* BCG-Danish strain was provided by Dr. Marcel Behr (RI-MUHC, Montreal). All strains were grown either in liquid 7H9 medium (Difco) supplemented with 10% ADC, 0.2% glycerol (Sigma-Aldrich) and 0.05% Tween-80 (Sigma-Aldrich), or on 7H11 agar (Difco) plates supplemented with 10% OADC and 0.5% glycerol. The antibiotics RIF, INH and KAN were purchased from Sigma-Aldrich, whilst AMK and CAP were purchased from Cayman Chemical Company (supplied by Cedarlane).

### DNA purification, PCR and sequencing

*Mtb* genomic DNA was isolated and purified according to the protocol of Pelicic *et al*. (21). For PCR-based screening of BCG clones picked into 7H9/ADC, boiled culture lysates were prepared (*e.g*. 250μl of culture heated at 90°C for 30mins, followed by centrifugation and resuspension of the pelleted material in 50μl TE). Taq DNA Polymerase, 10X reaction buffer, MgCl_2_ and dNTPs were obtained from Thermo Fisher Scientific. PCR was carried out according to standard protocols except for when amplifying a portion of the KAN resistance cassette where 5% DMSO was also included in the reaction mixtures. Primers used in this study for PCR and sequencing are shown in Supplementary Table S1, a number of which are based on those reported by Rowneki *et al.* (22). Sanger sequencing of PCR products was carried out at the Centre d’expertise et de services Génome Québec (Montreal).

### Minimal inhibitory concentration (MIC) determination

Two-fold serial dilutions of antibiotics were added to 96-well microtitre plates prior to the addition of an equal volume of *Mtb* or BCG culture diluted 1:100 from growing stock cultures adjusted to an OD_600nm_ of 0.1. The plates were sealed in plastic zip-lock bags and incubated for 7-14 days at 37°C. 30μl of 0.01% Resazurin (Sigma-Aldrich) was added to each well and the plates incubated for a further 4 days prior to quantifying fluorescence on a Tecan Infinite 200 Pro plate reader. All MIC assays were set up in triplicate or quadruplicate, with independent assays repeated on at least two occasions.

### Mycobacterial recombineering

Oligo-mediated recombineering was carried out essentially as described by Murphy *et al*. (19). Briefly, BCG-Danish was transformed with the pNitET-SacB-kan plasmid (referred to from hereon as pNitET; available from Addgene) and selected on 7H11/OADC agar plates containing 20μg/ml KAN. A single BCG::pNitET positive clone was subsequently grown to an OD_600nm_ of approximately 0.8 and treated for 24hrs with 1μM isovaleronitrile (Sigma-Aldrich) to induce expression of the RecET proteins prior to co-transformation of the 70bp oligos (Invitrogen; Table S1) targeting *rpoB* (0.1μg) and *rrs* (1μg). A control (no DNA) electroporation was also included. The electroporated cells were allowed to recover in 10ml of 7H9/ADC medium at 37°C with shaking for 4 days, following which a 1ml aliquot was diluted 1:20 with 7H9/ADC containing STR at 1μg/ml. Once growth of the antibiotic treated cultures was detected and they had reached an OD_600nm_ of approx. 0.3 (after 15 days), 150μl aliquots of undiluted and diluted (1:10) cells were spread onto 7H11/OADC agar plates containing either 1μg/ml STR, 2μg/ml RIF, or both. No KAN was added to the media. After 4 weeks of incubation at 37°C, colonies were picked into 1ml 7H9/ADC without antibiotic (to further aid in curing the bacteria of the pNitET plasmid) and allowed to grow for 3 weeks with occasional mixing. A sample of the cultures were screened for the presence of the desired *rrs* mutation by sequencing of PCR products generated from boiled culture lysates. The complete loss of the pNitET plasmid from the clones of interest was also confirmed by PCR using primers specific to the KAN resistance cassette.

## RESULTS

### MIC analysis and sequence determination

Initial drug susceptibility testing (DST) by the LSPQ classified the Lineage 4 Montreal *Mtb* isolate #57001 as being resistant to RIF, INH and STR. To more accurately assess the MICs of these compounds towards this strain, we conducted broth microdilution assays in 96-well format. In this manner, we determined the MICs to be 250μg/ml for RIF, 0.2μg/ml for INH and 4μg/ml for STR, which served to confirm the initial report of the local Public Health laboratory. Then, to identify the genetic basis for resistance towards each of these antibiotics, PCR products corresponding to the genes that are most frequently associated with mutations conferring resistance towards these compounds were amplified and sequenced. This included products corresponding to the 81bp rifampicin resistance determining region (RRDR) of *rpoB,* the *katG, inhA* and *inhA* promoter sequences (associated with INH resistance), as well as the *rpsL, rrs* and *gidB* genes, mutations in which are most frequently responsible for resistance towards STR (10, 22). In this manner we identified that the strain possessed the S450W RpoB mutation that is frequently linked to RIF resistance, and the −15C/T *inhA* promoter mutation consistent with the low-level of INH resistance observed for strain #57001. Far more surprising was the finding that the strain lacked any of the commonly reported *rpsL*, *rrs* or *gidB* alleles frequently associated with resistance towards STR. The only variation from the wild-type (H37Rv) sequence we could identify in any of these sequences was a G to A SNP at position 878 (G878A) of the *rrs* 16s rRNA gene. This corresponds to nucleotide 1472729 of the complete NCBI H37Rv reference sequence (NC_018143.2; 2012 release).

A BLAST^®^ search of the NCBI nucleotide database was also somewhat surprising in that whilst we were able to identify additional strains bearing the identical *rrs* G878A SNP, there appeared to be only 4 of them within the database. Likewise, a search of the literature identified relatively few publications making reference to *Mtb* strains bearing the G878A mutation. What was also striking was that there was no obvious consensus amongst these articles regarding exactly which antibiotic resistance profile the G878A mutation might be associated with. As discussed below, these articles variably referred to the strain(s) bearing this mutation in the context of either AMK, CAP, KAN or STR, and there was certainly no experimental confirmation of these largely genomics-based epidemiological studies (14–18).

### Introduction of the G878A *rrs* mutation into wild-type antibiotic-sensitive *M. bovis* BCG-Danish via oligo-mediated recombineering

In an effort to clarify the role of the G878A *rrs* SNP in aminoglycoside resistance, we decided to precisely engineer this mutation within the chromosome of a non-pathogenic Mycobacterial species that is very closely related to *Mtb*, namely *M. bovis* BCG (Danish str.). Prior to this we re-confirmed that the BCG-Danish strain was fully susceptible to the antibiotic compounds of interest for this study (*i.e*. AMK, CAP, KAN & STR; Table 1). 70bp oligos targeting the relevant portions of the *rrs* and *rpoB* genes and containing the desired G878A and C1349T SNPs at their centre were co-introduced into the BCG-Danish background via electroporation at a ratio of 10:1, respectively. The *rpoB* oligo was originally designed for use as a recombineering control as it encodes the well characterized S450L substitution commonly associated with acquired RIF resistance. However, its use turned out to be quite fortuitous as it allowed us to select for our recombineered clones that had taken up both oligos (STR/RIF resistant) in amongst the large background of spontaneous STR mutants that appeared following growth and selection at the relatively low concentration of 1μg/ml STR. Note that this amount of STR was chosen based on the MIC value observed with *Mtb* isolate #57001, *i.e*. 4μg/ml. We reasoned that spontaneous mutants - independent of the recombineering process - were also being selected for at 1μg/ml STR due to the fact that we observed equivalent growth and CFU (colony forming units) following plating of the no DNA control, as well as for an additional control in which a 70bp oligo containing the wild-type *rrs* sequence was used in place of that containing the G878A SNP (also delivered in conjunction with the oligo bearing the mutant *rpoB* allele). Sanger sequencing of PCR products spanning the targeted *rrs* sequence for a selection of 12 clones isolated following co-transformation of the *rrs*^G878A^ + *rpoB*^C1349T^ oligos and plating at 1μg/ml STR, confirmed that mutations other than the G878A SNP were being selected for in this case. Additional sequencing revealed that all 12 clones contained one of four distinct *gidB* polymorphisms: W45*(stop), 352insG (insertion), 468insA or 532insG. Notably, a broad range of *gidB* mutations have previously been associated with low-level STR resistance in *Mtb*, which is consistent with the apparent ease of their *in vitro* selection herein at 1μg/ul STR (10, 23).

**Table 1:** Representative *in vitro* MIC values for the aminoglycosides AMK, CAP, KAN & STR.

|  | **AMK<sup>\*</sup>** | **CAP** | **KAN** | **STR** |
| --- | --- | --- | --- | --- |
| **BCG-Danish** | 0.08 | 0.31, 0.4<sup>#</sup> | 0.8, 1.6<sup>†</sup> | **0.16** |
| **BCG-RpoB S450L** | 0.08 | 0.31 | 0.8 | **0.16** |
| **BCG-rrs G878A/**<br>**RpoB S450L** | 0.08 | 0.16, 0.2<sup>#</sup> | 0.8, 1.6<sup>†</sup> | **1.25** |

<sup>\*</sup> All MIC values are presented as µg/ml.
<sup>#</sup> A separate dilution series was used in this case leading to a slightly different MIC value.
<sup>†</sup> Different values were obtained in independent assays.

After plating the original post-transformation outgrowth cultures (enriched at 1μg/ml STR) onto 7H11/OADC plates supplemented with either 2μg/ml RIF or 1μg/ml STR + 2μg/ml RIF, colonies were only obtained for the transformation that included both the G878A *rrs* and C1349T *rpoB* containing oligos. Neither of the control transformations (no DNA & wild-type *rrs* oligo controls) resulted in even a single colony in this case. Five clones from each of the successful platings were sequence confirmed to have incorporated the two expected SNPs within their *rrs* and *rpoB* genes, respectively. A single clone from each independent plating (BCG-Danish_*rrs*^G878A^/*rpoB*^C1349T^ clones #1 & #15) was then selected for downstream analysis after confirming via PCR that they had been cured of the pNitET (KAN^R^) recombineering plasmid. In addition, we also sequence confirmed that these clones did not carry any additional, unwanted mutations within their *gidB* or *rrs* genes that could have confounded our subsequent analyses.

### Introduction of the G878A *rrs* mutation results in resistance to STR, but not other relevant aminoglycosides used in the treatment of TB

Broth microdilution assays in 96-well format and incorporating two-fold serial dilutions of AMK, CAP, KAN & STR were used to examine the relative impact of the G878A *rrs* mutation on the MICs obtained for the two recombineered clones (#1 & #15) in comparison to the parental BCG-Danish strain. To control for any unforeseen effect that introduction of the S450L RpoB mutation may have had on the MICs obtained in the presence of these aminoglycoside antibiotics, two RIF resistant clones selected from the mutant *rpoB*/ wild-type *rrs* oligo (control) transformation at 2μg/ml RIF were also tested in MIC assays relative to the BCG-Danish wild-type. In this manner, we established that the S450L RpoB substitution had no measurable impact on the response towards AMK, CAP, KAN or STR (Table 1). Thus, we could be confident that any MIC alterations we observed when testing the recombineered *rrs* mutants were solely the result of introducing the G878A SNP.

As shown in Table 1, introduction of the G878A SNP resulted in a reproducible 8-fold increase in the MIC for streptomycin with respect to the wild-type and RIF-resistant controls (0.16 vs. 1.25μg/ml). On each occasion that they were tested, both the mutant clones (#1 & #15) behaved identically in these assays. As the MIC was raised above the critical breakpoint concentration reported by the WHO for STR in broth culture (MGIT culture; 1μg/ml) (24) this result confirms our hypothesis that the G878A *rrs* SNP confers acquired resistance towards STR. However, we see no evidence that this SNP confers cross-resistance to any of the other major aminoglycosides used in the treatment of antibiotic resistant TB: AMK, CAP or KAN. If anything, we noted what appears to be a minor (2-fold) increase in susceptibility towards CAP (Table 1).

### Introduction of the G878A *rrs* mutation into BCG-Danish impacts fitness when grown in the absence of antibiotic

Although the G878A mutation clearly confers low-level STR resistance, we were curious as to why we had some difficulty in obtaining recombineering clones bearing this mutation in the absence of the secondary selection step in the presence of RIF. We reasoned that the mutation may be deleterious to the strain in some manner that impacts its relative growth and fitness. To examine this hypothesis, we carried out standard *in vitro* growth curves in liquid 7H9/ADC broth to compare the relative growth of the G878A mutant clones (#1 & #15) to the parental BCG-Danish strain and the *rpoB* mutation control in the absence of added antibiotic. As can be seen from Fig. 1(a), introduction of the G878A SNP into *rrs* leads to a substantial reduction in growth rate, above-and-beyond that observed with the S450L RpoB mutation alone that is commonly associated with RIF resistance in *Mtb*. Over the first 96 hours, where the growth rates were mostly linear for all the strains examined, the relative doubling times were as follows: BCG-Danish wild-type (28.7 hours); BCG-Danish_*rpoB*^C1349T^ (32.3 hours; average for clones #2 & #3); BCG-Danish_*rrs*^G878A^*rpoB*^C1349T^ (39.9 hours; average for clones #1 & #15).

**Figure 1.**
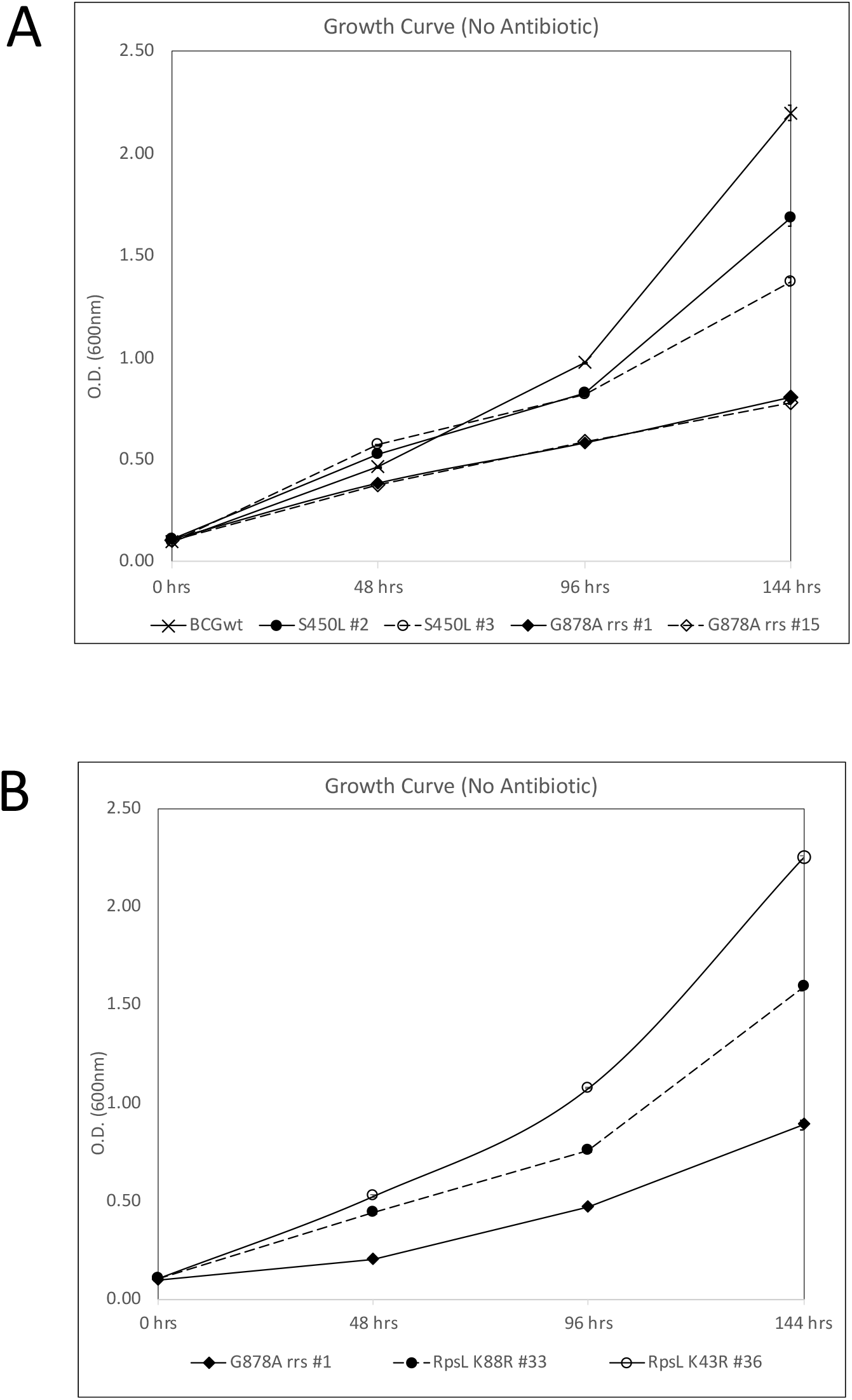
Introduction of the *rrs* G878A mutation results in a growth/ fitness defect *in vitro.* **A)** Growth curves carried out in 7H9/ADC media (in the absence of added antibiotic) comparing wild-type *M. bovis* BCG-Danish against two RIF-resistant (BCG-Danish_*rpoB*^C1349T^; S450L clones #2, #3) and two STR/RIF resistant (BCG-Danish_*rrs*^G878A^*rpoB*^C1349T^; G878A *rrs* clones #1, #15) clones obtained by recombineering. Two separate OD_600nm_ readings were obtained for all 5 cultures at each of the time points indicated. Standard deviations are indicated by crosshairs. For the wild-type and both of the G878A *rrs* mutant clones, the data are representative of two independent growth assays. **B)** Growth curves carried out in 7H9/ADC media (no antibiotic) comparing STR/RIF-resistant strains carrying three distinct combinations of mutations: BCG-Danish_*rrs*^G878A^*rpoB*^C1349T^ (G878A *rrs* clone #1), BCG-Danish_*rpoB*^C1349T^_*rpsL*^A263G^ (RpsL K88R #33), and BCG-Danish_*rpoB*C1349T_*rpsL*A128G (RpsL K43R #36). As above, two separate OD_600nm_ readings were obtained for all cultures at each of the time points indicated and the data are representative of two independent growth assays. Standard deviations are indicated by crosshairs.

In a separate assay, we then compared BCG-Danish_*rrs*^G878A^*rpoB*^C1349T^ clone #1 against two RIF/STR-resistant clones that were generated by selecting the BCG-Danish_*rpoB*^C1349T^ (S450L) RIF-resistant mutant on 2μg/ml STR. These clones were identified as carrying the archetypal STR resistance-conferring mutations, RpsL K88R (clone #33) and RpsL K43R (clone #36). As shown in Fig. 1(b), the recombineered strain bearing the *rrs*^G878A^ SNP shows a clear growth/ fitness defect relative to both of these RpsL mutant strains. Although not directly compared “head-to-head” within the same growth assay, by comparing the curves shown in Fig. 1, we noted that introduction of the RpsL^K43R^ mutation seemingly improved the growth rate of the RpoB^S450L^/RpsL^K43R^ double mutant relative to that of the RIF-resistant RpoB^S450L^ single mutant. The RpsL K43R mutation is by far the most common mutation associated with STR resistance in clinical *Mtb* isolates, particularly amongst MDR isolates, which is consistent with its low fitness-cost (10, 25, 26). In addition, Spies *et al*. have previously noted that amongst a small sample of MDR/STR-resistant isolates, those with the RpsL K43R allele showed growth enhancement, even relative to fully drug-susceptible *Mtb* isolates (27). Indeed, this type of unexpected epistatic interaction whereby dual-resistant strains bearing particular combinations of mutations show improved fitness over the corresponding mono-resistant strains has been described previously in relation to RIF and ofloxacin resistance (28). Nevertheless, as it was not the main objective of the current study, we have not investigated this phenomenon in relation to the RpoB S450L and RpsL K43R alleles any further at this stage.

In summary, our data support the hypotheses that the G878A *rrs* mutation results in a substantial fitness defect - at least under standard *in vitro* conditions in the absence of antibiotic. However, at this point we cannot exclude the possibility that the fitness defect attributed to the G878A *rrs* mutation may only manifest itself in the context of strains also bearing the S450L RpoB mutation (or similar).

## Discussion

Although the G878A *rrs* mutation that is the focus of this investigation appears to be relatively uncommon amongst antibiotic resistant patient isolates, one of the primary motivations for our study was to clarify which, if any, of the second-line aminoglycoside antibiotics it conferred resistance towards. We felt that clarification was necessary in this case due to the potential confusion that could arise based on surveying the currently available literature. For example, of the 5 manuscripts we identified as reporting the G878A SNP, 3 associated its presence with isolates that were STR resistant (15, 17, 18). Only one of these studies looked at resistance to other aminoglycosides in addition to STR (CAP, KAN), but did not find an association with the G878A SNP (15). Curiously, we also found 2 studies that examined the G878A *rrs* SNP in the context of aminoglycosides other than STR. In one of these studies, a single isolate bearing the G878A polymorphism was shown to be susceptible to AMK, CAP and KAN (14). Then, a South-African study of pre-XDR and XDR-TB patients identified 21 isolates with the G878A mutation (16). However, the distribution of the mutation with respect to the reported resistance phenotypes was quite variable: 10 of the isolates were CAP mono-resistant, 4 were KAN mono-resistant, 4 were cross-resistant to AMK/CAP/KAN, 2 were AMK/CAP cross-resistant, and the remaining isolate was resistant to CAP/KAN. As per the previous study, none of the isolates were examined for STR resistance for reasons that were not given. Based on the frequency of CAP resistance, the G878A *rrs* SNP was reported by these authors as a new mechanism of resistance towards CAP. As such, it was recommended that the G878A mutation be included in new molecular assays to increase the sensitivity of CAP resistance detection (16). Finally, in addition to the 4 isolates identified by BLAST^®^ searching of the NCBI database, we have also identified 4 distinct MDR-TB isolates included within the PATRIC (Pathosystems Resource Integration Center) database that are reported to carry the G878A *rrs* allele (29). Three of these are classified as being STR resistant, whilst all four are listed as susceptible to AMK, CAP and KAN. Overall, we felt it quite important to investigate this polymorphism at the experimental level in an attempt to generate conclusive data directed at addressing the nagging question regarding its precise clinical relevance.

Our initial identification of the G878A *rrs* mutation was in the context of an MDR-TB isolate, that along with RIF and INH resistance was classified by the Provincial public health laboratory as being STR resistant. We subsequently confirmed that the isolate exhibited low-level STR resistance (approx. 4μg/ml), which immediately suggested that it did not contain either of the polymorphisms within the *rpsL*-encoded S12 ribosomal protein (K43R, K88R) that are commonly associated with high-level STR resistance (>32μg/ml) (10, 23, 30). Through sequencing, we also ruled out the possibility that the isolate carried any of the *gidB* (encodes a 16s rRNA methyltransferase) alleles that tend to be seen in isolates with low-level STR resistance (10, 23, 25), nor did this *Mtb* isolate possess any other *rrs* mutation aside from the G878A polymorphism. Although there are relatively few published studies where we find the G878A *rrs* mutation mentioned, it is interesting to note that in each case the *Mtb* isolates involved are either MDR or XDR. This is despite the fact that STR mono-resistance is second only to INH mono-resistance in terms of global frequency (25). Whether this observation reflects some form of cryptic epigenetic interaction between specific mutations in *rpoB* and the *rrs* G878A mutation, for example, that lead to a measurable enhancement of *in vivo* fitness over *Mtb* cells bearing the *rrs* G878A mutation alone, or whether it reflects a strong bias towards the detection, sequencing and reporting of MDR/ XDR-TB clinical isolates at present rather than those that are mono-resistant, is not clear. Alternatively, it may reflect the reality that STR has been relegated to use only in second-line regimens due to problems of resistance and patient toxicity (ototoxicity and nephrotoxicity). We do note, however, that our engineered BCG-Danish_*rrs*^G878A^*rpoB*^C1349T^ strain exhibits a substantial fitness defect *in vitro* whereby its observed doubling time is 1.24X that of the *rpoB*^C1349T^ RIF-resistant mutant strain. Nevertheless, this is not necessarily reflective of the situation *in vivo,* including when antibiotics are being applied, nor does it account for the possibility that additional compensatory mutations may also evolve during a natural infection process that may, at least in part, mitigate the negative consequences of acquiring and maintaining this mutation. It does suggest, however, that in the absence of any compensatory adaptation, cells that arise with the G878A *rrs* polymorphism are likely to be at a distinct competitive disadvantage in the presence of other resistant clones that do not exhibit a fitness defect to the same degree. The *in vitro* growth defect we report may also be relevant to the clinical microbiology laboratory’s ability to detect or isolate cells having the G878A SNP from patient samples, particularly in the context of heteroresistance (*i.e*. mixed populations of resistant *Mtb* isolates). As such, the G878A *rrs* SNP may simply be underreported. Either of these scenarios might well explain why *Mtb* isolates containing the G878A mutant *rrs* allele are quite rare amongst clinical isolates appearing in the NCBI and PATRIC databases, as well as in the published TB literature.

In terms of an underlying mechanistic basis that could potentially explain how the G878A rrs SNP may contribute to resistance towards STR, we note that position 878 of the *Mtb* 16s rRNA molecule is equivalent to position 885 within a highly-conserved region of the *Escherichia coli* 16s rRNA sequence (31–35). In *E. coli*, residue 885 (G) basepairs with residue 912 (C) at the base of helix 27, a structure implicated in tRNA selection in both prokaryotes and eukaryotes (36). Cross-linking, footprinting and mutagenesis experiments have all demonstrated that STR binds to this same area - specifically to residues 912 - 915 that form what is referred to as the “915 region” (31, 37, 38). Moreover, at least two independent studies have shown that mutations introduced into this 915 region - including a C to T mutation at position 912 - reduces STR binding to the ribosome resulting in low-level resistance towards this antibiotic (31, 39, 40), analogous to the phenotype we report herein for both the *Mtb* isolate #57001 and our recombineered BCG-Danish_*rrs*^G878A^*rpoB*^C1349T^ strain. By inference, we hypothesize that a G to A substitution at position 878 within the *Mtb* or BCG *rrs* sequence will prevent its base pairing to the complementary cytosine residue at position 905 (equivalent to *E. coli* residue 912). In turn, this disruption has the potential to perturb the organization of the 915 region in a manner that may impede the binding of STR, thereby leading to resistance (Figure 2). Notably, each of the other 3 aminoglycosides examined in this study, namely AMK, CAP & KAN, have all been shown to bind to a distinct region of the 16s rRNA molecule known as the “A-site” (aminoacyl-tRNA site) that comprises a portion of helix 44 (41, 42). Thus, mutations causing resistance towards these compounds all tend to be localized around *rrs* position 1400 (10, 23, 25).

**Figure 2.**
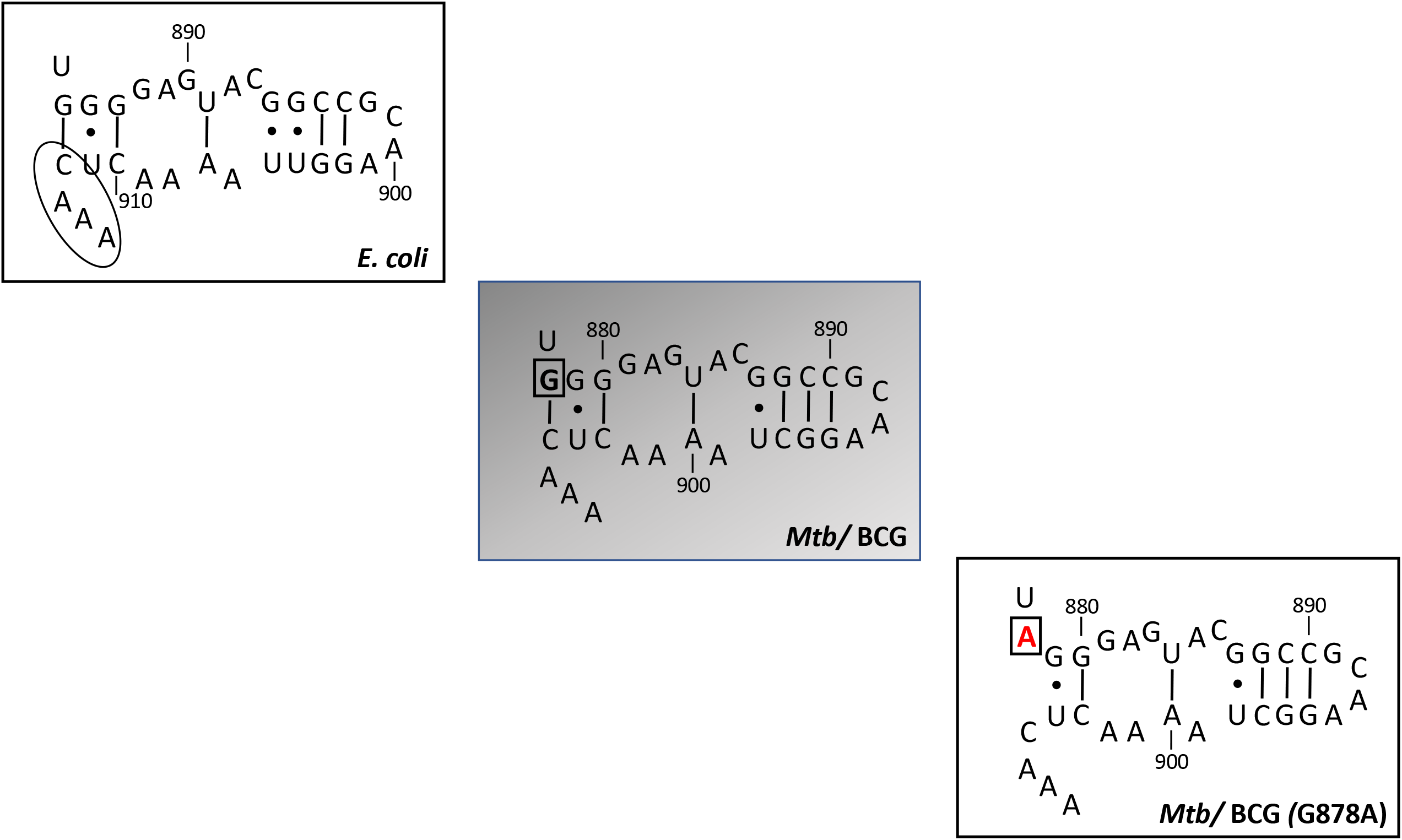
Predicted structure of the 16s rRNA helix 27 region in *Mtb* and *M. bovis* BCG based on that of *E. coli* (top panel). Note that the *E. coli* and *Mtb*/ BCG sequences are identical in this region aside from a single C substitution at position 897 in the latter. Canonical basepairs within the helix are indicated by lines, while “wobble” (G-U) base pairing is indicated by dots. The position of the G878A *rrs* mutation is indicated in red (bottom panel). The relative nucleotide numbers for both the *E. coli* and *Mtb*/BCG 16sRNA sequences are included at 10bp intervals (900 etc.). The 915 region involved in STR binding is circled within the *E. coli* panel. These schematic representations are adapted from Cannone *et al*. (2002) (35) and the RiboVision2 website [http://apollo.chemistry.gatech.edu/RiboVision/index.html] (34).

In summary, through a combination of genetic recombineering and *in vitro* MIC assays, herein we have - for the first time - experimentally confirmed that the presence of the clinically relevant *rrs*^G878A^ mutation causes low-level STR resistance. However, by itself, it does not alter susceptibility to the other second-line injectable aminoglycosides used in the treatment of TB. In addition to providing an important point of clarification regarding the precise role of this SNP, this knowledge is also relevant in light of recent calls for the reinstatement of STR for use in the treatment of drug-resistant TB caused by isolates that exhibit low-level STR resistance, yet are highly resistant to AMK and KAN (23).

