## Supplemental Table S1 for "Experimental confirmation that an uncommon, yet clinically relevant mutation (G878A) in the *rrs* gene of *Mycobacterium tuberculosis* confers resistance to streptomycin"

| Oligos for Mutation Screening (PCR + Sequencing): | | | | | Product Size (bp) | Notes |
|---|---|---|---|---|---|---|
| ***rpoB* RRDR** | rpoB_RRDR_F | CGATCACACCGCAGACGTTG | rpoB_RRDR_Rev | CACGCTCACGTGACAGACC | 170 | |
| ***katG*** | katG_F | GGCGGTCACACTTTCGGTAA | katG_R | CTGTTGTCCCATTTCGTCGG | 191 | Rowneki *et al*. 2020 PLoS ONE |
| ***inhA*** | inhA_F | CACAAGGACGCACATGACAGG | inhA_R | TCCTCGAGCAGCTGGATCTG | 672 | |
| ***inhA* promoter** | INHprom_F | CCCAGAAAGGGATCCGTCAT | INHprom_R | GATACGAATGGGGGTTTGGC | 204 | Rowneki *et al*. 2020 PLoS ONE |
| ***rpsL*** | rpsL_F | GGTCAAGACCGCGGCTCTGA | rpsL_R | GTAGACCGGGTCGTTGACCA | 375 | Rowneki *et al*. 2020 PLoS ONE |
| ***rrs*** | rrs_F | TCGGATTGACGGTAGGTGGA | rrs_R | CATTCCACCGCTACACCAGG | 225 | Rowneki *et al*. 2020 PLoS ONE |
| | | | rrs_R2 | CTTCCGGTACGGCTACCTTG | 1049 | Used with rrs_F for PCR |
| | rrs_F2 | GTCCAGGGCTTCACACATGC | | | | Used for sequencing |
| | | | rrs_R3 | CCTGCACACAGGCCACAAG | 575 | Used with rrs_F to confirm presence of G878A mutation in recombineering clones |
| **gidB** | gidB_F | CAGCGTCTCGAGAGCGGAG | gidB_R | CGTCGGTGTCGGTGGTGTC | 852 | |
| **Kanamycin** | KanF | ATGAGCCATATTCAACGGGAAA | KanR | CAAACCGTTATTCATTCGTGATTG | 480 | To confirm loss of pNitET; used with 5% DMSO |

| Recombineering Oligos (mutations in bold and underlined): | |
|---|---|
| **rrs G878A** | TTCCTTTGAGTTTTAGCCTTGCGGCCGTACTCCC**T**AGGCGGGGTACTTAATGCGTTAGCTACGGCACGGA |
| **rrs wild-type** | TTCCTTTGAGTTTTAGCCTTGCGGCCGTACTCCCCAGGCGGGGTACTTAATGCGTTAGCTACGGCACGGA |
| **rpoB S450L** | CACGCTCACGTGACAGACCGCCGGGCCCCAGCGCC**A**ACAGTCGGCGCTTGTGGGTCAACCCCGACAGCGG |

**Table S1: Oligonucleotides used in this study for PCR, sequencing and recombineering.**
